# Lipid membrane templated misfolding and self-assembly of intrinsically disordered tau protein

**DOI:** 10.1101/2020.03.04.977421

**Authors:** Jaroslaw Majewski, Emmalee M. Jones, Crystal M. Vander Zanden, Jacek Biernat, Eckhard Mandelkow, Eva Y. Chi

**Author notes:** Name of Corresponding Author: Eva Y. Chi.

## Abstract

The aggregation of the intrinsically disordered tau protein into highly ordered β-sheet fibrils is implicated in many neurodegenerative disorders. Fibrillation mechanism remains unresolved, particularly early events that trigger tau misfolding and assembly. We investigated the role membrane plays in modulating aggregation of three tau variants, the largest isoform hTau40, the truncated construct K18, and a hyperphosphorylation mutant hTau40/3Epi. Despite being charged and soluble, tau proteins were also highly surface active and favorably interacted with anionic, but not zwitterionic, lipid monolayer at the air/water interface. Membrane binding induced macroscopic tau phase separation and β-sheet-rich tau oligomer formation. Concomitantly, membrane morphology and lipid packing became disrupted. Our findings support a general tau aggregation mechanism wherein tau’s inherent surface activity and favorable electrostatic interactions drive tau-membrane association, inducing tau phase separation that is accompanied by misfolding and self-assembly of disordered tau into β-sheet-rich oligomers, which subsequently seed fibrillation and deposition into diseased tissues.

## Introduction

Intrinsically disordered proteins (IDPs) comprise a class of proteins that defy the structure-function paradigm as they do not have well-defined 3D structures. Rather, they occupy a structural ensemble of rapidly inter-converting conformations, allowing IDPs to carry out important functions during which they often undergo structural transitions^1–6^. It is also recognized that structural plasticity of IDPs is in part responsible for their disproportional prevalence in diseases, including neurodegenerative disorders, cancers, and cardiovascular diseases^7–17^. The misfolding and aggregation of tau, a microtubule-associated IDP, into highly ordered β-sheet-rich fibrils, paired helical filaments (PHFs) that subsequently deposit into neurofibrillary tangles (NFTs) inside neurons are implicated in a range of neurodegenerative disorders, tauopathies, that include Alzheimer’s disease (AD), Pick’s disease, and frontotemporal dementia with parkinsonism-17^13, 18^. Despite our increased understanding tau physiology and pathology^19–21^, a key feature of tau pathology, tau aggregation, remains to be resolved at the structural and mechanistic level.

The aggregation of natively folded proteins has been found to involve at least two critical steps. The first requires the perturbation of the protein’s native structure to form an aggregation-competent intermediate that proceeds through the formation of a structurally expanded transition state^22–24^. The second step involves the assembly of intermediates into aggregates. The energetics of the two steps are respectively governed by the conformational stability of the protein native state and colloidal stability of the protein solution^25, 26^. In contrast to natively folded proteins, IDPs have very little conformational stability and the disordered native state is often stable and resistant to aggregation. Native tau is highly soluble, contains many charged and hydrophilic residues, and shows little tendency for aggregation^12, 21, 27^. Thus, tau aggregation must proceed through a different mechanism and requires a different type of structural perturbation to render tau aggregation-competent.

Tau aggregation involves the transition of a disordered monomeric state into a highly ordered fibrillar state (Fig. 1). In contrast to folded proteins, this ordered aggregation^28^ pathway first involves the formation of a partially folded aggregation-competent intermediate (A_I_) and proceeds through a structurally contracted transition state (N^‡^) (Fig. 1). A_I_ then irreversibly assemble into higher order aggregates (A_m_). A high energy barrier for tau partial folding, i.e., large Δ*G*^‡^ stemming from entropic penalty, therefore, results in slow tau aggregation. Conditions that favor compact conformations of tau or induce disordered-to-ordered transitions can thus reduce Δ*G*^‡^ and drive aggregation.

**Figure 1:**
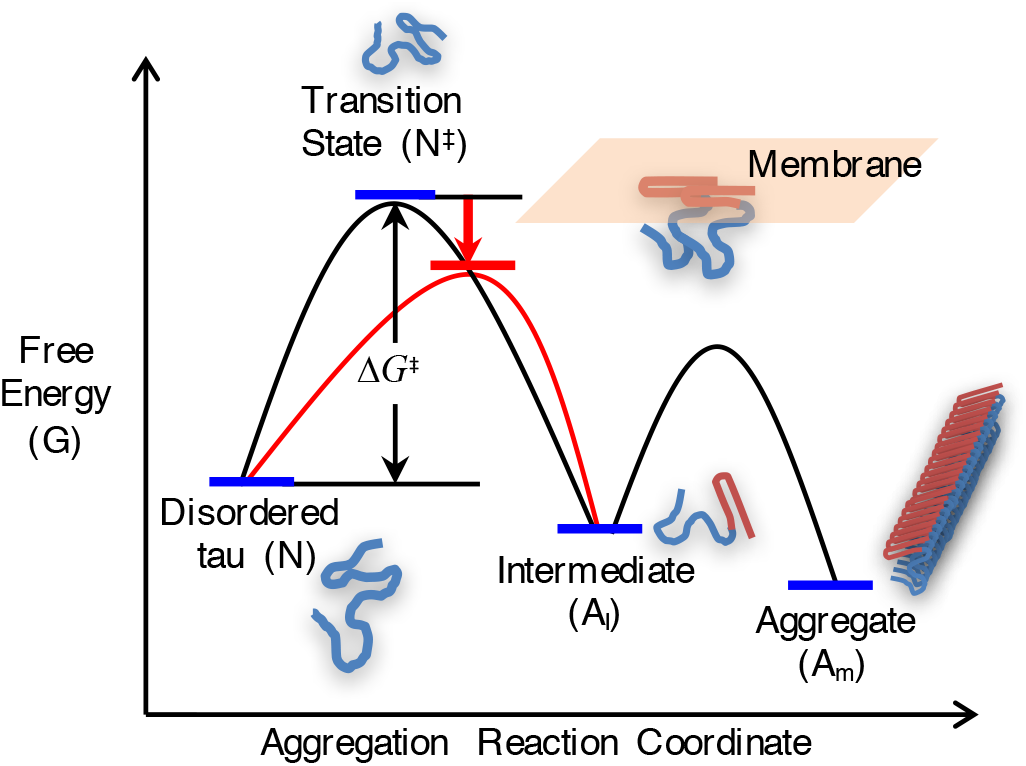
Schematic of tau aggregation pathway from a disordered native state (N) to a highly ordered fibrillar aggregate state (A_m_) that proceeds through the formation of a partially folded, aggregation-competent intermediate (A_I_). Conditions that lower the free energy of a structurally compacted transition state (N^‡^), such as favorable tau-membrane interactions, can induce disordered-to-ordered transitions, lowering the activation free energy (Δ*G*^‡^) of aggregation and accelerate fibril formation (modified from Chi et al. ^25^).

*In vitro* studies have shown that tau aggregation follows a heterogeneous nucleation-elongation pathway where aggregation is accelerated by polyanionic cofactors, including heparin, RNA, and arachidonic acid micelles^27, 29–34^. More recently, a few studies have shown that tau undergoes liquid-liquid phase separation (LLPS) and that the phase-separated tau can serve to initiate tau aggregation^35–37^. Anionic lipid membranes have also been shown to efficiently facilitate tau aggregation^27, 31, 38, 39^ and therefore provide a physiologically relevant context for tau aggregation *in vivo*. Tau has been shown to favorably interact with anionic membranes^40–43^, although details of the structural changes and assembly associated with membrane binding that leads to fibril nucleation remain to be elucidated. We previously studied the interaction between the full-length wildtype tau, hTau40, with zwitterionic and anionic lipid monolayers at the air/water interface ^44^. Our results showed that although highly soluble and charged, hTau40 is also highly surface active and preferentially interacted with anionic lipid membrane. Moreover, hTau40 associated with anionic lipid membrane is more conformationally compact compared to its disordered state in solution, indicating a possible membrane-mediated aggregation pathway for tau assembly (46) (Fig. 1).

To evaluate if the surface and membrane interactions observed for hTau40 are a general property of tau, we investigated the interactions between three different tau proteins (Fig. 2) with anionic and zwitterionic lipid membranes and used *in situ* high resolution synchrotron X-ray scattering methods to resolve structural details of tau-membrane interactions. The wildtype full length hTau40, truncated isoform K18, as well as a hyperphosphorylation mutant (hTau40/Epi) were used to explore the effect of tau domain composition and hyperphosphorylation on its surface activity and membrane interactions. K18 (residues Q244-E372) lacks the N-terminal projection domain and the C-terminal tail, but contains the repeat domains (repeats R1-R4) that form the core of tau fibrils ^45^. Phosphorylation of tau is developmentally regulated^46^, and in tauopathies, tau is abnormally hyperphosphorylated, particularly at or near the flanking domains^47^. Phosphorylation in the repeat domain (e.g. in the KxGS motifs) reduces tau affinity to microtubules and triggers detachment from microtubule surface. The interplay between different phosphorylation sites, tau conformation, and interactions is still poorly understood. In this study, we used a pseudo-hyperphosphorylation mutant of hTau40, hTau40/3Epi, wherein 7 serine and threonine residues in the flanking domains were mutated into anionic glutamic acid residues (Fig. 2). These sites correspond to the epitopes of 3 antibodies (AT8, AT100, PHF1) that recognize phospho-tau at early stages of neurodegeneration. Although glutamic acid is not a perfect substitute for phosphorylation, it is a reasonable approximation and has the advantage of being controllable compared to incomplete and heterogeneous phosphorylation by various kinases^48^.

**Figure 2:**
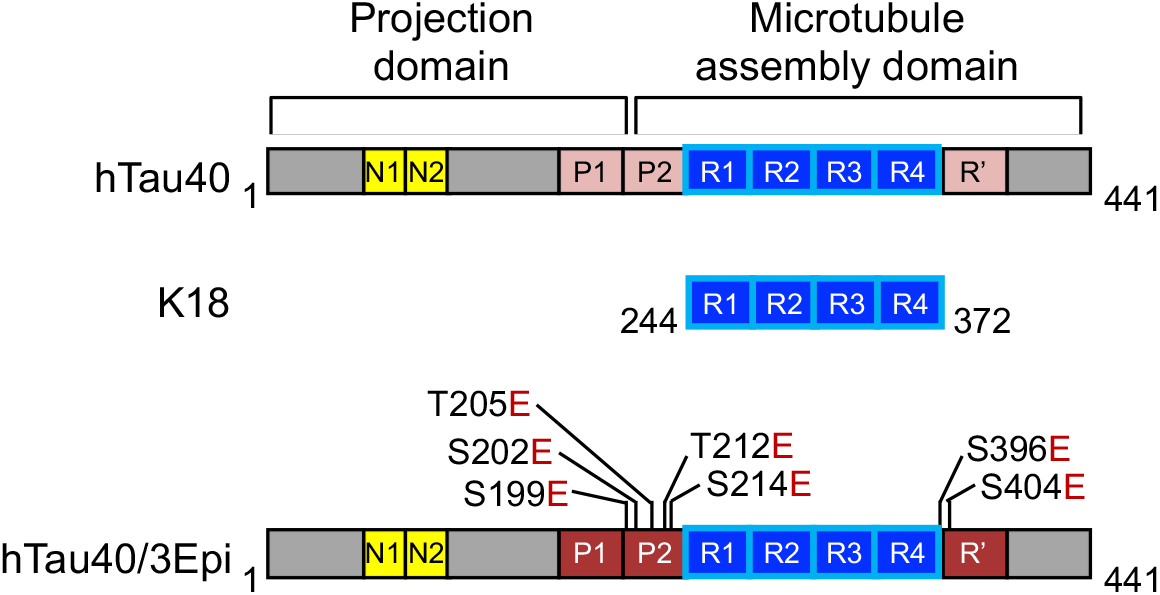
Schematics of domain compositions of the three tau proteins studied. Wildtype hTau40 (441 residues) is the largest human isoform in the central nervous system. Tau construct K18 (Q244-E372) contain the 4 repeats (R1-R4) and hTau40/3Epi is a pseudo-hyperphosphorylation mutant with 7 extra anionic residues. The serine (S)/threonine (T) to glutamic acid (E) mutations cover three key epitopes of antibodies against phospho-tau which require 2 (or 3) adjacent phosphorylation sites, recognized by the epitopes AT8 (residues S199, S202, T205), AT100 (T212, S214), and PHF1 (S396, S404). These epitopes are used as diagnostic markers for incipient neurodegeneration. Net charges on tau proteins are: hTau40 = +2, K18 = +10, and hTau40/3Epi = −5.

Interactions between the three tau proteins with lipid monolayers at the air/water interface were measured. *In situ* synchrotron grazing incidence X-ray diffraction (GIXD) was used to resolve angstrom-level structural details of tau and membrane at the air/water interface. Our results show that anionic membranes induced macroscopic tau phase separation, and templated the misfolding and assembly of tau into β-sheet-rich oligomers, providing direct evidence of membrane-templated tau misfolding and aggregation.

## Results

### Tau proteins are highly surface and membrane active

We have previously shown that although highly charged and soluble, the full-length hTau40 is highly surface and membrane active^44^. To evaluate if these properties are intrinsic to the protein, we assessed the surface and membrane activities of K18 and hTau40/3Epi proteins. Surface activity was evaluated by measuring surface pressure (π) isotherms of tau adsorption to the air/water interface from a water subphase in a Langmuir trough where π is the magnitude by which the surface tension (γ) of a clean air/water interface (γ_o_) is reduced by the presence of an adsorbate (π = γ_o_ − γ) such as proteins or lipids. The air/water interface serves in both experimental and computational studies as a model hydrophobic/hydrophilic interface to mimic the cell’s organic/aqueous interfaces. In this study, adsorption of tau to the air/water interface was characterized to assess tau’s propensity interact with a purely hydrophobic surface, which contributes significantly to its interactions with more complex bio-interfaces, such as membranes.

All tau proteins readily adsorbed to the air/water interface from the bulk causing immediate and rapid increases in π followed by more gradual increases (Fig. 3A). These features reflect different stages of tau adsorption wherein diffusion and adsorption of the protein from the bulk to the interface caused the initial rapid increases in π. Structural rearrangement of the adsorbed tau as well as addition of new tau to the adsorbed layer caused further and slower increases in π. Adsorption of the mutant hTau40/3Epi is moderately slower, although the final π reached was slightly higher than that of hTau40. The 7 extra anionic residues of the mutant, which changed tau’s net charge from +2 to −5, increased repulsive protein-protein electrostatic interactions and may have contributed to the reduced adsorption rate, but did not alter tau’s accumulation at the interface. The shorter K18 construct also readily adsorbed to the air/water interface, but at an even slower rate and reached a lower π value (Fig. 3A). K18 has a net charge of +10 and the strong repulsive protein-protein interactions likely caused the slow adsorption rate as well as the lower π reached. Nonetheless, our results show that the MTB domains of tau, which forms the core of PHFs, is surface active and contributes significantly to the overall surface activity of tau.

**Figure 3:**
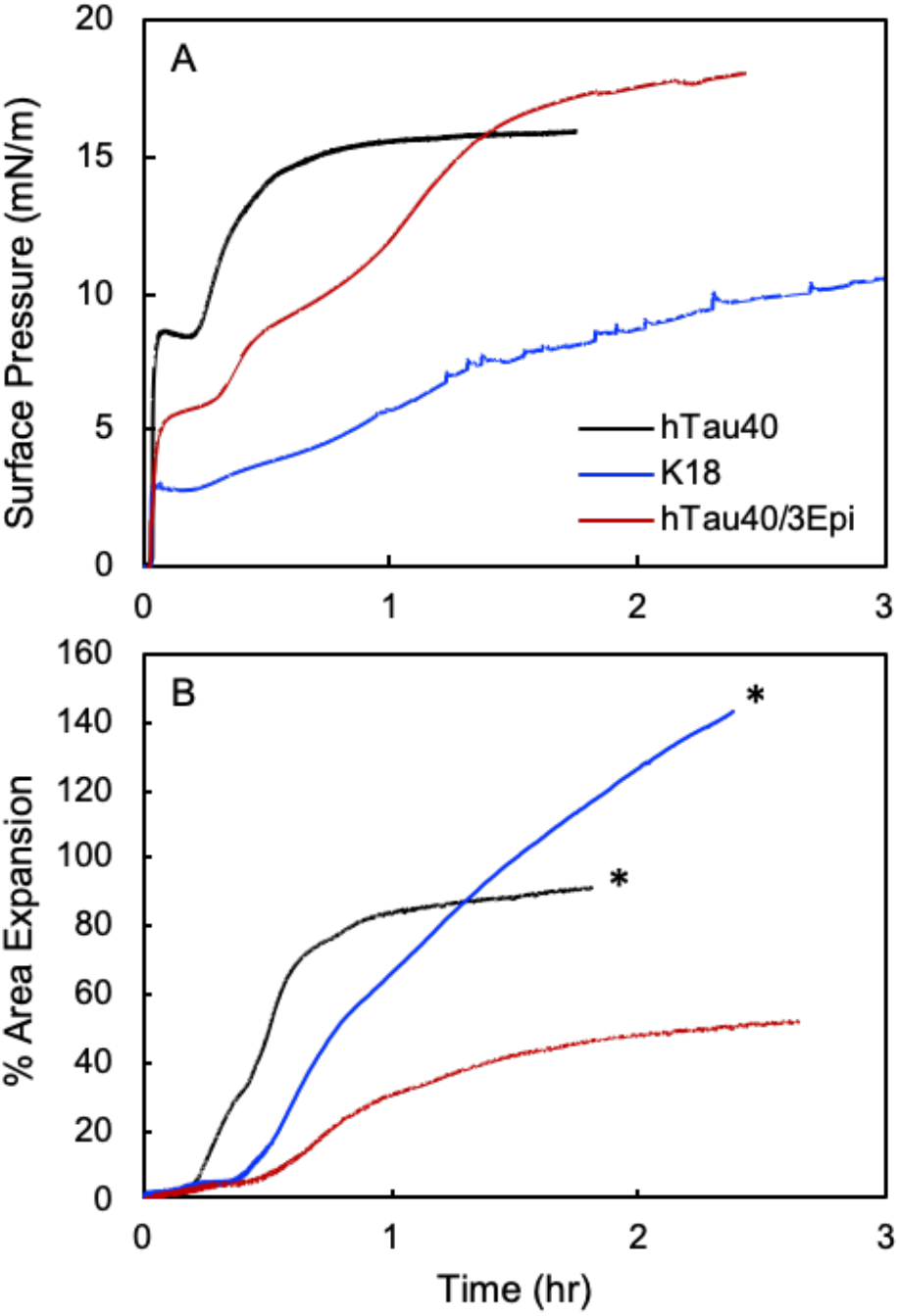
**(A)**: Adsorption isotherms of 1 μM tau to the air/water interface at 25°C. Time 0 corresponds to the time of tau injection into the water subphase of a Langmuir trough. Surface pressure (π) is defined as π = γ_o_ − γ, where γ_o_ is the surface tension of a clean air/water interface and γ the surface tension of the air/water interface with an adsorbate. **(B)**: % area expansion profiles of 1 μM tau inserting into DMPG monolayers held at 25 mN/m on water subphase at 25°C. A lipid monolayer was first compressed to 25 mN/m, after which tau was injected into the subphase (time = 0). Insertion of tau into the membrane due to favorable tau-lipid interactions result in expansion of the monolayer surface area. In two experiments marked by *, the lipid monolayers became completely expanded in the Langmuir trough and further expansion was not measurable.

Many factors influence protein surface activity. Non-polar residues contribute to hydrophobicity and are expected to strongly favor protein adsorption and influence orientation at the air/water interface. Moreover, as multi-point contacts between the protein and the surface are usually required for favorable adsorption^49, 50^, conformational flexibility plays an important role in a protein’s surface activity. Structurally flexible, or “soft” proteins readily undergo structural rearrangements at an interface to make more contacts. For natively folded proteins, this type of structural changes usually results in protein denaturation upon surface adsorption. For tau, the disordered native state allows the protein to easily rearrange to maximize desolvation and expose non-polar residues to the air phase. Our results indicate that this driving force is sufficiently strong so that increases in tau-tau repulsion from the 7 extra anionic residues at phosphorylation sites did not significantly impact tau’s surface activity. It is also clear that both the projection domain as well as the MTB domains contribute to tau’s surface activity as the adsorption of the two larger tau proteins resulted in larger π values compared to K18. Overall, our results indicate that tau’s surface activity is an intrinsic property of the “soft” protein, and one that is not significantly affected by variations in domain or amino acid composition.

In addition to being surface active, all tau proteins were also membrane active (Fig. 3B). Despite their different charges, each tau protein inserted into an anionic 1,2-dimyristoyl-*sn*-glycero-3-phospho-(1’-rac-glycerol) (DMPG) lipid monolayer held at a constant π of 25 mN/m, causing monolayer area expansions that ranged from 50% to over 140%. This π value was chosen for its relevance to physiological conditions as the lipid packing density of a bilayer membrane has been reported to correspond to that of a monolayer at around 25-30 mN/m ^51^. In contrast, tau proteins did not insert into zwitterionic 1,2-dipalmitoyl-sn-glycero-3-phosphocholine (DPPC) lipid monolayers at 25 mN/m (data not shown), indicating no favorable interactions with the membrane. Compared to cationic hTau40, the anionic hTau40/3Epi inserted into the DMPG monolayer at a slower rate and caused about half the area expansion. (Fig. 3B)

Favorable interactions between K18 and the related K19 construct to anionic lipid membranes have been attributed to favorable protein-lipid headgroup electrostatic interactions^40, 42^. Our results show that that despite the overall anionic nature of hTau40/3Epi, the hyperpho-mutant readily and favorably interacted with the anionic membrane, demonstrating a strong driving force for tau membrane binding that overcomes electrostatic repulsions. Not surprisingly, the strongly cationic K18 construct caused significantly larger monolayer area expansion compared to hTau40. We note that due to experimental constraints, the insertion of hTau40/3Epi and K18 did not reach final equilibrated states, where the Langmuir trough barriers became fully expanded during these experiments. Thus, further protein insertion, especially for K18, was not measured. Nonetheless, it is clear from our results that although hyperphosphorylation reduced tau-membrane interactions, it did not prevent the protein from interacting with and intercalating into the anionic membrane. Truncation of the tau projection domain significantly enhanced the interactions of the MTB domains with DMPG membrane as K18 was the most membrane active.

### Tau proteins disrupt membrane morphology

To assess changes in lipid monolayer morphology caused by the insertion of the tau proteins, fluorescence microscope (FM) images of the lipid monolayer were taken before and at various time points after the injection of tau during constant-π insertion assays. Representative images are shown in Fig. 4 and percent area expansion is indicated for each image. At 25 °C, the DMPG monolayer on water subphase undergoes a liquid-expanded (LE) to lipid-condensed (LC) phase transition at around 17 mN/m^52^. Because the bulky lipid dye molecules, TR-DHPE (0.5 mol%), are preferentially excluded from the LC phase, it appears as dark domains whereas the fluid LE phase containing the dye is bright.

**Figure 4.**
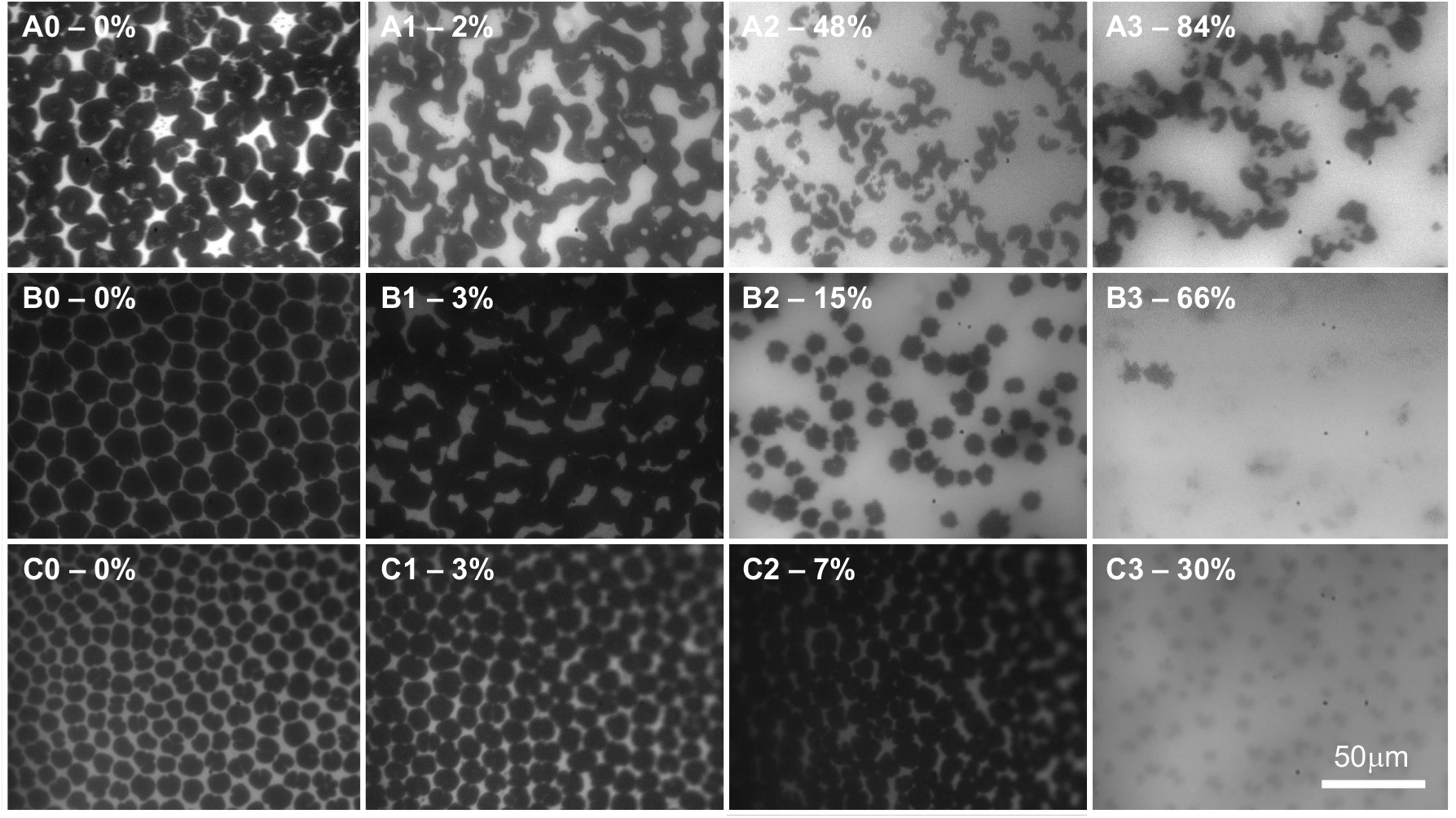
Fluorescence microscopy (FM) images of DMPG lipid monolayers at 25 mN/m before (0) and 10 (1), 30 (2), and 60 (3) minutes after injecting hTau40 (**A**), K18 (**B**) or hTau40/3Epi (**C**) into the subphase. % insertion of tau at the time points are also indicated. Head group-labeled fluorescent dye TR-DHPE (0.5 mol%) was included in the lipid monolayer to provide fluorescence contrast. Dark regions of the monolayer correspond to dye-excluded lipid-condensed (LC) domains whereas light, or fluorescent region of the monolayer corresponds to disordered liquid-expanded (LE) phase of the monolayer where the TR-DHPE dye is mixed with the DMPG lipids.

As shown, DMPG monolayer at 25 mN/m contained predominantly dark LC phase domains (Fig. 4A0, 4B0, and 4C0). At 10 min after protein injection where minimal protein insertion had occurred (Fig. 4A1, 4B1, 4C1), the LC domains changed from a well-defined and circular shape to less well-defined shapes where many of the LC domains had fused. These changes indicate decreased line tension around the LC domains due to the association of tau to the lipid monolayer but before any significant tau insertion had occurred. At 30 minutes after injection, where significant insertion had occurred for hTau40 and K18, 48% and 66%, respectively, the morphology of the monolayer changed drastically (Fig. 4A2 and 4B2). The ratio of dark to light regions was reduced and the LC domains became smaller, especially for hTau40. These changes indicate that tau insertion into the disordered phase had pushed the LC domains apart, as well as disrupted the LC phase. hTau40/3Epi inserted into the membrane at a slower rate and caused about 7% area expansion 30 minutes after injection. Although the monolayer contained primarily LC domains, the domains fused and became less defined (Fig. 4C2), similar to those observed for hTau40 and K18 insertion at the 10 min time point with low insertion levels (Fig. 4A1 and 4B1). At 1-hour post injection, the morphology of the lipid monolayer incubated with hTau40 (Fig. 4A3) did not significantly differ from that at 30 min post injection, although area expansion almost doubled during that time period. However, morphologies of monolayers from K18 and hTau40/3Epi insertion significantly changed. The LC domains became almost completely disrupted where distinct domains boundaries were no longer present.

The FM images of the lipid monolayer indicate that the association of the tau proteins to the lipid monolayer caused observable changes to the monolayer morphology even in the absence of significant insertion at earlier time points. Subsequent insertion of hTau40 primarily occurred in the fluid LE phase of the monolayer. The LC phase domains were perturbed with hTau40 insertion, but the domains retained their distinct boundaries. However, insertion of the more highly charged proteins K18 and hTau40/3Epi, caused almost complete disruption of the LC phase, indicating that these two proteins were more membrane disruptive than hTau40.

### Anionic membrane templates misfolding and assembly of tau into β-sheet-rich aggregates

Using synchrotron GIXD, we monitored the time-dependent *in situ* insertion and accumulation of the tau proteins with lipid DPPC and DMPG monolayers. A lipid monolayer at 25 mN/m was first formed and an aliquot of tau was then injected into the water subphase underneath the monolayer. In these experiments, as the Langmuir trough surface area was held constant, insertion of tau into the monolayer caused increases in π; final π values ranging from 34 to 46 mN/m were obtained for the different tau proteins. GIXD measurements were collected before (*t* = 0) and at approximately 2.5 and 12 hours after tau injection. As shown previously, at 25°C DMPG monolayer contained ordered LC phase giving rise to in-plane *Bragg peaks* which allowed us to track changes to lipid packing during tau insertion.

In GIXD experiments, the X-ray beam strikes the surface at an incident angle below the critical scattering angle for total external reflection. At this angle, an X-ray evanescent wave is generated and penetrates a few nanometers into the bulk liquid^53^. The wave travels along the surface and Bragg scatters from two dimensional (2D) ordered molecular arrangements at the air/water interface. GIXD measurements therefore provide in-plane (*i.e.* in the plane of the monolayer) structural information on the diffracting, or ordered, portion of the film. In our experiments, the lipid alkyl tails in LC phase and ordered tau assemblies give rise to diffraction peaks. The reciprocal space GIXD patterns from the 2D ordered structures (*Bragg peaks* and *rods*) can be analyzed to provide complete structural information about molecular arrangement of lipids and the interacting proteins^54–57^.

Fig. 5 shows GIXD data obtained for DMPG monolayer before (Fig. 5A) and after injecting each tau in the subphase (Fig. 5B-5D). Structural parameters extracted from the diffraction peaks are summarized in Tables S1, S2, and S3 in Supplemental Information. Fig. 5A shows two *Bragg peaks* resulting from the LC phase of the DMPG monolayer, indicating a distorted hexagonal 2D arrangement of the alkyl tails^57^. The broader peak at lower *Q*_xy_ is the superposition of two {1,0}+{0,1} Bragg reflections and the sharper peak at higher *Q*_xy_ is the {−1,1} reflection. Peak broadness is inversely proportional to the average size of the 2D ordered domain, or coherence length *L*_xy_^58^, which are 170 Å and 310 Å for the {1,0}+{0,1} and {−1,1} directions, respectively (Table S1B). hTau40 caused significant changes to the lipid packing (Fig. 5B1 and 5B2). The intensity of the lipid diffraction peaks decreased where *L*_xy_ values were 100 and 240 Å for the {1,0}+{0,1} and {−1,1} directions, respectively, and the integrated intensity reduced by 87% after 12 hours of tau incubation (Fig. 5B2 and 6B, Table S1). Strikingly, decrease in lipid packing was accompanied by the appearance of a new diffraction peak marked with a red asterisk at *Q*_xy_ = 1.34 Å^−1^. This *Q*_xy_ value corresponds to a *d*-spacing of 4.74 ± 0.02 Å, which exactly matches the average distance between β-sheet arrangements in tau fibrils^59^. *L*_xy_ of the β-sheet crystallites was around 200 Å (Table S1C), indicating that there were approximately 42 tau β-strands arranged in positional registry. Our results thus show that the insertion of hTau40 into DMPG monolayer induced two concomitant structural changes, tau-induced disruption of lipid-packing and membrane induced assembly of the disordered tau protein into ordered β-sheet aggregates.

**Figure 5:**
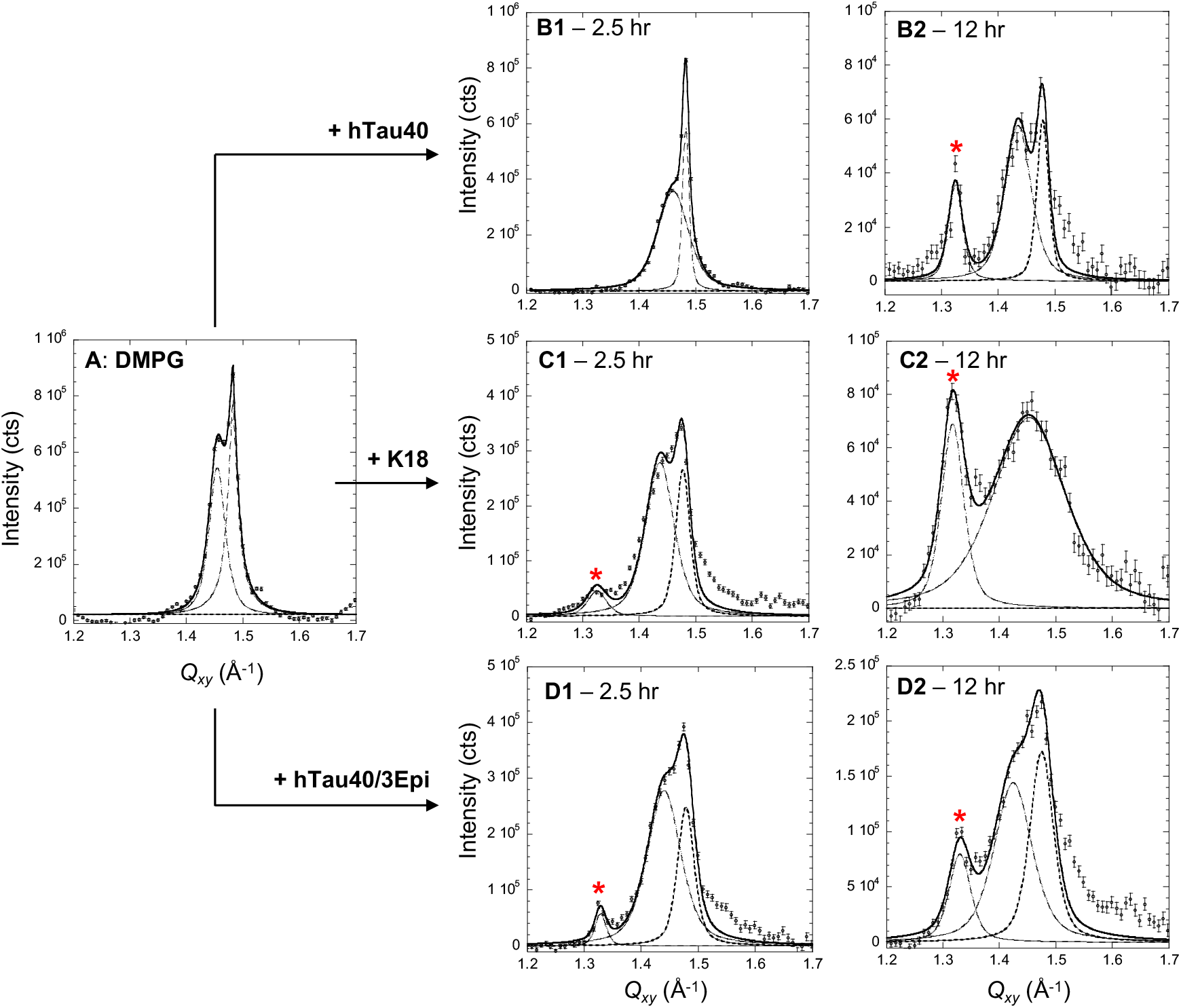
*Bragg peaks* from DMPG monolayer at the air/water interface at 25 mN/m and 25°C before **(A)**and *t*_1_ = 2.5 and *t*_2_ = 12 hours after the injection of hTau40 **(B)**, K18 **(C)** and hTau40/3Epi **(D)** proteins. The *Bragg peaks* were fitted using the sum of two Voigt profiles (solid line) and de-convoluted into separate peaks (dashed lines) corresponding to {1,0}+{0,1} and {1,−1} *Bragg peaks* in the distorted hexagonal 2D unit cell. *Bragg peaks* were obtained by integrating over the −0.05 Å^−1^ ≤ *Q_z_* ≤ 0.75 Å^−1^ region. Red asterisks indicate *Bragg peaks* associated with ordered β-sheet structures of the tau protein. Structural parameters extracted from GIXD and *Bragg rods* are presented in Supplemental Information.

**Figure 6:**
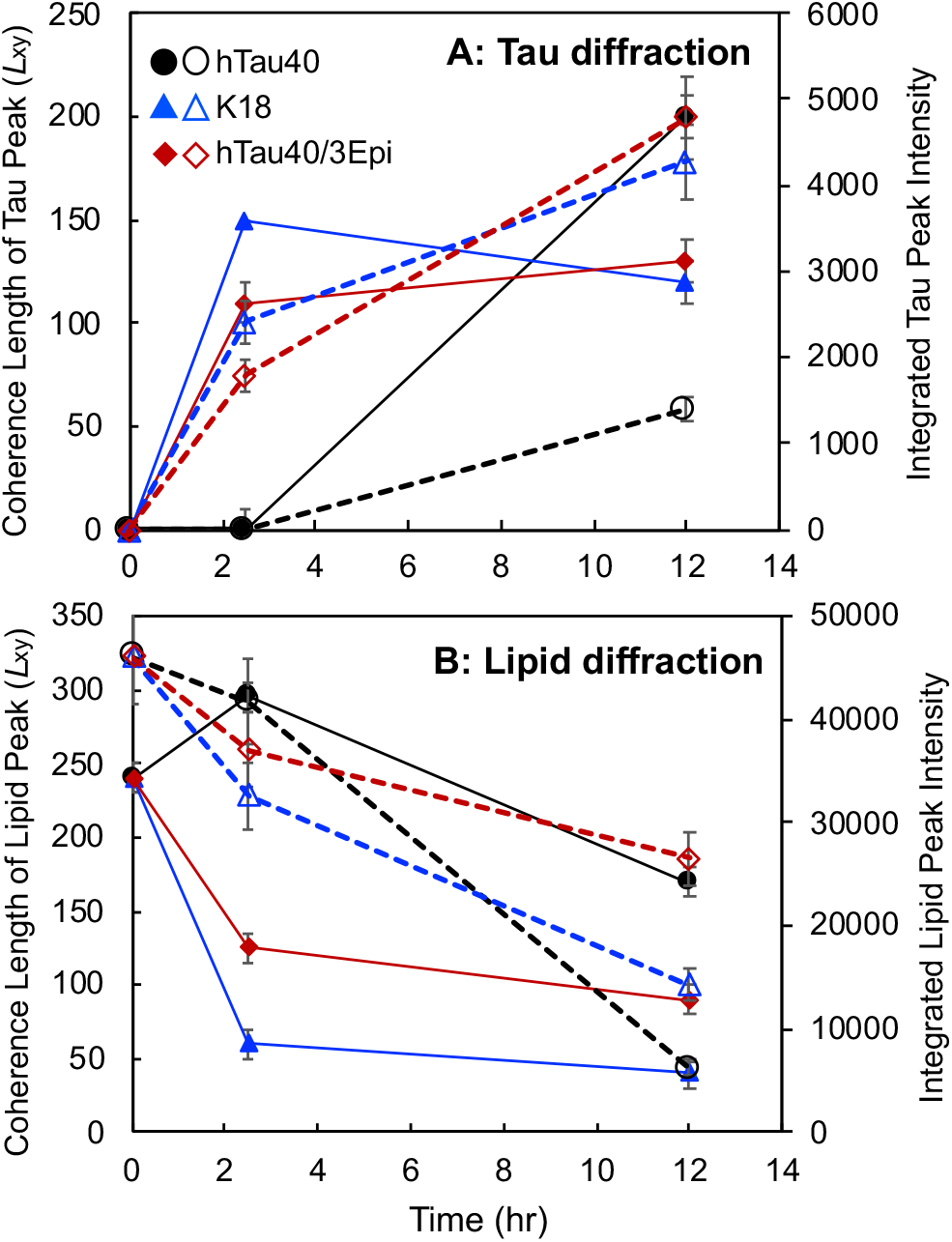
Coherence length (*L*_xy_ in Å) (filled symbols and solid lines) and integrated intensities (unfilled symbols and dashed lines) of the tau *Bragg peaks Q*_xy_ = 1.34 Å^−1^ (**A**) and lipid *Bragg peaks* (**B**) extracted from GIXD measurements of tau insertion into DMPG monolayers. For lipids, averaged *L*_xy_ values of the {1,0}+{0,1} and {1,−1} *Bragg peaks* were plotted.

GIXD results for K18 and hTau40/3Epi interacting with DMPG monolayers also showed the two salient features of hTau40-membrane binding, membrane-templated β-sheet-rich tau aggregate formation and tau-induced membrane disruption (Figs. 5C-5D and 6). However, important differences were also observed. K18 caused rapid decreases in the intensity of the lipid diffraction peaks where the two original peaks converged into one broad reflection 12 hours after K18 injection (Fig. 5C1 and 5C1), which had a *L*_xy_ value of only 40 Å. The integrated intensities of the lipid peaks reduced to 70% and 30% of the original value at 2.5 and 12 hours, respectively, after K18 injection. These results indicate that the LC domains became more disrupted with K18 insertion compared to hTau40. This result is consistent with FM images where K18 caused the most disruption to LC domains at the morphological scale. The remaining ordered structures corresponded to short-ranged hexagonal packing of lipid tails with increased unit cell vectors from 4.93 to 5.00 Å, indicating an increase in area per lipid molecule in the monolayer, even as the π of the film was significantly higher, at 46 mN/m. Importantly, the β-sheet ordered tau aggregates appeared earlier for K18 than hTau40, where the *Bragg peak* at *Q*_xy_ = 1.34 Å^−1^ was already visible at 2.5 hr. After 12 hours of incubation, *L*_xy_ value of the ordered K18 peak did not significantly change but the intensity of this peak increased two-fold, indicating that while size of the ordered K18 aggregates did not grow, their number doubled from 2.5 to 12 hours of incubation (Fig. 5C1-5C2 and 6, Table S2).

Similar to K18, hTau40/3Epi caused significant reductions to the amount of ordered LC phase in the DMPG monolayer and gave rise to in-plane β-sheet diffraction peak (marked by asterisk) (Fig. 5D1 and 5D2). Despite the strong repulsive protein-protein and protein-membrane electrostatic interactions, the mutant formed the tau diffraction peak earlier compared to hTau40. By 2.5 hours of incubation (Fig. 5D1), the tau diffraction peak was visible and the amount of tau aggregates grew by almost three-fold by 12 hours (Figs. 5D and 6, Table S3).

In addition to analyzing the *Bragg peaks*, the analysis of their *Bragg rods* was also carried out to obtain the length of the coherently scattering moieties participating in the Bragg reflection (*L*_c_). For lipid scattering, *L*_c_ is the length of the coherently scattering portions of the alkyl tails measured along their backbones. For protein scattering, *L*_c_ is the length of the coherently scattering β-sheet portion of the protein. Although the *Bragg rod* data are in general noisy (Figs. S1-S3 in Supplemental Information), we were able to extract *L*_c_ values for the protein scattering peaks and they ranged from 4.5 – 7.2 Å. The dimension of the β-sheet strand backbone has been reported to be approximately 10 Å^59^. Our analysis thus indicates that the β-sheet moiety that coherently participate in the Bragg reflection is one layer of β-sheets.

In contrast, ordered aggregates were not observed for tau incubated with zwitterionic DPPC monolayers (data not shown). The in-plane structure of the DPPC monolayer also remained unchanged in the presence of tau proteins (data not shown). These results show that a lack of protein-membrane interactions resulted in a lack of structural changes to both the protein and membrane. GIXD data were also measured for the tau films adsorbed at the air/water interface and no in-plane diffraction peak was observed (data not shown). Concentrating tau at the air/water interface is thus insufficient to induce β-sheet formation. Favorable interactions between tau and anionic lipids are thus necessary to promote tau β-sheet formation and aggregation.

## Discussion

Taken together, our results yielded molecular level details into the structure and dynamics of tau association with anionic membranes. As illustrated in Fig. 7A, hTau40 binding and insertion into the lipid membrane resulted in the structural compaction^44^ and misfolding of the disordered tau into aggregates wherein the protein is arranged in an extended β-sheet conformation that is one layer thick. K18 and the hyperphosphorylation mutant similarly formed β-sheet-rich aggregates at the membrane surface, but they formed earlier and larger numbers of aggregates were formed (Fig. S4 and S5). Concomitant to tau binding, lipid packing in the membrane became disrupted where the size of LC phase domains significantly decreased (Figs. 7D and 7E). Consistent with membrane morphology visualized by FM, K18 and hTau40/3Epi disrupted lipid packing in the membrane to greater extents (Figs. S4B, S4C, S5B, and S5C) compared to hTau40, implicating possibly higher levels of toxicity of these two tau proteins.

**Figure 7:**
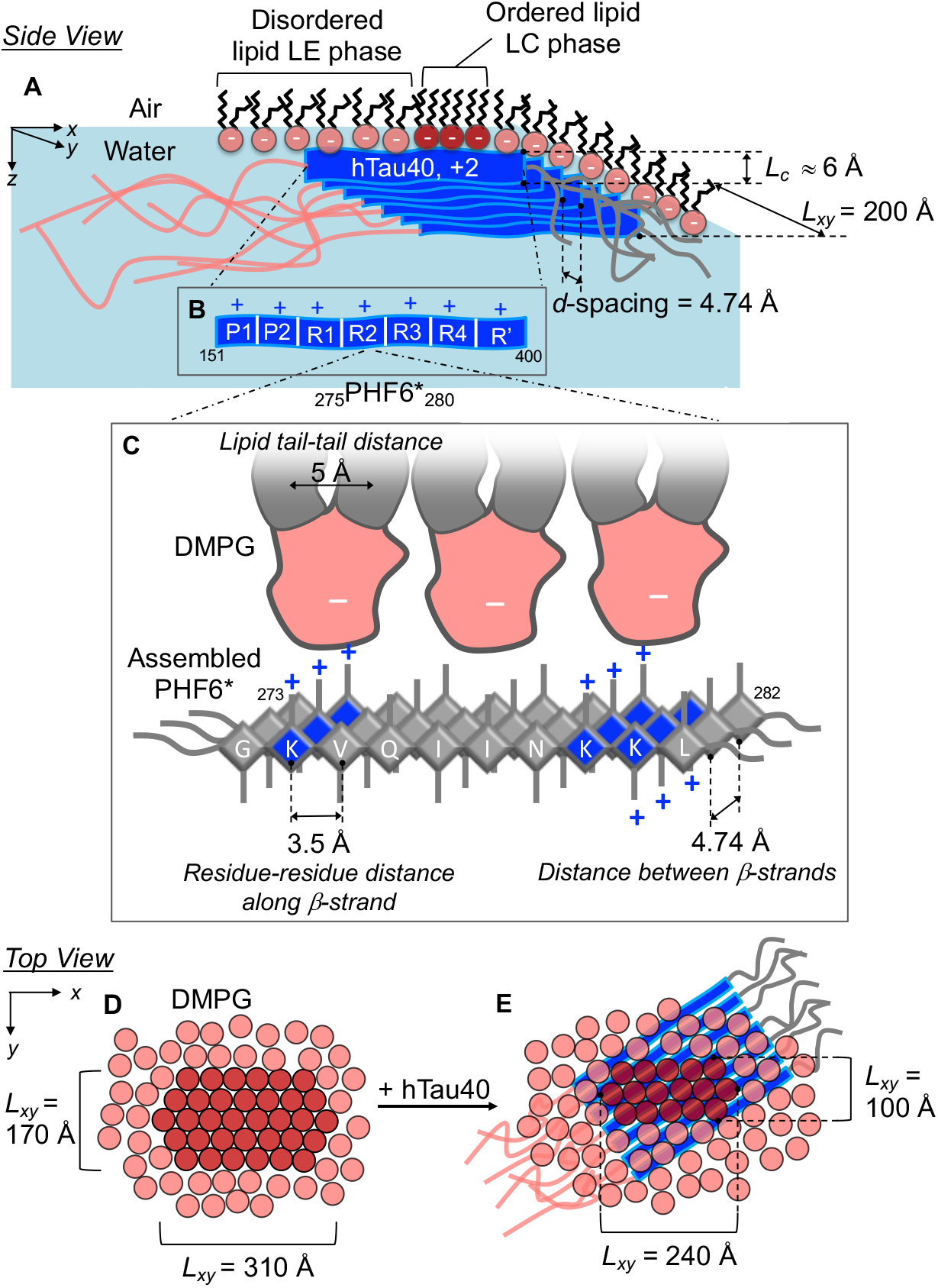
Schematics of a β-sheet enriched hTau40 oligomer templated by the anionic DMPG monolayer. **A-C**: side views (in the plane of the membrane) shows an ordered tau assembly associated with the anionic DMPG monolayer at the air/water interface (**A**). *d*-spacing and coherence lengths (*L_xy_*) shown are for the hTau40 oligomer. *L_c_* is the out-of-plane length of the coherently scattering moiety of the hTau40 assembly. Successive zoomed in schematics illustrate possible tau domains (cationic P1, P2 and P3) (**B**) and cationic amino acids flanking the hexapeptide PHF6* in the R2 domain that favorably interact with the membrane through attractive electrostatic interactions (**C**). **D-E**: top views (perpendicular to the membrane) of the membrane before (**D**) and after (**E**) hTau40 association. The *L_xy_* values in these schematics are those of the ordered lipid domains. The orientation of the tau β-sheet arrangements vis-à-vis the orientation of the DMPG lattice cannot be determined from the experimental data. Schematics for K18 and hTau40/3Epi can be found in Figures S5 and S6 in Supplemental Information, respectively.

The remarkable observation that all tau proteins formed β-sheet rich aggregates at the anionic membrane surface confirms results from earlier studies that showed nucleation of tau fibrils is strongly enhanced by lipid membrane^27, 31, 38, 39^. For the positively charged hTau40 and K18, our results show that the anionic membrane enhances protein-protein interactions by charge screening and concentrating the proteins at the membrane surface^40, 42^. Although the structural methods utilized in this study do not give domain or residue-specific information about tau-lipid binding, we suspect that the cationic microbutule binding (MTB) and flanking domains (P1 through R’) of hTau40 interact with the anionic lipid headgroups while the anionic N-terminus and neutral C-terminus do not interact with the membrane (Fig. 7A). This domain interaction model is further supported by the observation that the MTB domains alone (K18) strongly interacts with and forms β-sheet-rich aggregates at the anionic membrane surface. The MTB domains play major roles in both tau’s physiological function and aggregation and form the core of PHFs. In particular, two lysine-containing hexapeptide segments, ^275^VQIINK^280^ (PHF6*) and ^306^VQIVYK^311^ (PHF6) located in the second and third repeat units of the MTB domain, respectively, have been identified as aggregation promoting β-sheet forming motifs^60–62^. Our finding that the β-sheet-rich aggregates are only one sheet thick indicates that the backbone of the β-sheet strands is positioned perpendicular to the lipid surface such that alternate side chains of residues in the β-strand point upwards toward the membrane. Fig. 7C illustrates how such an arrangement might promote favorable electrostatic interactions between cationic lysine (K) side chains around the PHF6* sequence and anionic lipids. As shown, the distance between two lysines with side chains pointing towards the membrane, K274 and K280, is about 20 Å. Given that the tail-tail distance of the lipids in the membrane is about 5 Å, approximately three lipids span the cationic lysine-to-lysine β-strand backbone distance.

The recent tau LLPS studies show that tau demixing into protein-rich droplets is facilitated by crowding agents^35–37^ and the additional presence of polyanions such as heparin and RNA promotes tau fibrillation in the droplets. As LLPS appears to be a fundamental mechanism for organizing intracellular space^63^, LLPS of tau can lead to subcellular regions of high local tau concentration, which combined with other aberrant events such as hyperphosphorylation or aggregation prone tau, may lead to tau aggregation^35^. It also appears that LLPS alone is not sufficient and also not a necessary condition for tau aggregation. Results from this study demonstrate that anionic lipid membrane can potentially serve both roles, concentrating tau at the membrane surfaces as well as providing specific lipid-protein interactions that templated the formation of extended tau β-sheet formation. In fact, it appears that tau incubated with an anionic lipid membrane overtime forms a macroscopic gelatinous material at the air/water interface (Video S1 in Supplemental Material). This feather is not exhibited by the that the lipid membrane itself, nor by an anionic lipid membrane incubated with the amyloid-β (Aβ) peptide that also formed β-sheet crystallites at the membrane surface^64, 65^ (Video S2 in Supplemental Information). Although details of the tau gel formation are not yet clear, it appears that tau phase separation can also be induced by anionic membranes at a concentration as low as 1 μM.

Our finding that the anionic DMPG membrane also templated β-sheet formation of the anionic hTau40/3Epi points to a complex role of the membrane in mediating protein-protein interactions and misfolding. In addition to electrostatic interactions and concentration effects (e.g., higher local protein concentrations at membranes or in phase-separated tau droplets^35^), interactions with the hydrophobic lipid tails could also play an important role at facilitating tau-tau interactions and β-sheet assembly.

In conclusion, despite being highly charged and soluble, the flexible nature of the tau proteins tested also causes them to be highly surface active and all tau proteins favorably interacted with lipid monolayers composed of anionic DMPG, but not zwitterionic DPPC, lipids at the air/water interface. The K18 MTB domain construct exhibited the strongest interaction with the DMPG membrane and that the hyperphosphorylated state of tau found in disease states, albeit with a net negative charge, retains this selective affinity to anionic lipid membrane. Membrane binding also appeared to induce tau phase separation at the membrane surface. Strikingly, all tau proteins binding to DMPG monolayers gave rise to in-plane protein diffraction peaks which correspond to one layer of about 40 β-strands in positional registry. Thus, the membrane templated the formation of β-sheet rich tau oligomers at the membrane surface. Compared to the wildtype hTau40, K18 and phospho-mutant hTau40/3Epi formed β-sheet oligomers earlier and at larger numbers.

As tau aggregation is believed to be driven by a transition from random coil to β-sheet structure, our study shows that the lipid membrane effectively catalyzed this structural transition for further growth into mature fibrils. In-plane protein diffraction peaks were not observed for tau incubated with zwitterionic DMPC lipids nor tau adsorbed at the air/water interface. Thus, the accumulation of the protein, or increasing protein local concentration, alone is insufficient to induce tau β-sheet formation. This is in contrast with our previous findings that the amyloid-β (1-40) peptide formed in-plane diffraction peaks both adsorbed at the air/water interface^66^ and inserted into anionic lipid membranes^65^. Although the exact nature of tau-membrane interactions that leads to tau structural re-organization and assembly into extended β-sheet structures remains to be resolved, the effect is membrane specific, but not tau domain composition-specific. Our findings support a general tau aggregation mechanism where tau’s inherent surface activity drives tau-lipid membrane interactions, inducing the misfolding and self-assembly of tau into β-sheet enriched oligomers that could subsequently seed tau fibril growth and deposition into diseased tissues. Concomitant with misfolding of the tau protein, tau-membrane interactions also results in the disruption of the lipid packing in the membrane, pointing to a possible membrane disruption-based toxicity pathway of tau oligomers.

## Methods

### Materials

hTau40, K18 and hTau40/3Epi were expressed and purified^48, 67, 68^. 1,2-dimyristoyl-sn-glycero-3-[phospho-rac-(1-glycerol)] (DMPG) and Dipalmitoylphosphatidylcholine (DPPC) were purchased from Avanti Polar Lipids (Alabaster, AL). Lipid stock solutions ranged from 2 to 10 mg/ml were prepared by dissolving lipids in chloroform (Fisher Scientific, Hampton, NH) containing 10 vol% methanol and then diluted to 0.2 or 0.5 mg/ml for spreading solutions. For fluorescence microscopy, the head group-labeled fluorescent dye Texas Red 1,2-dihexadecanoyl 3-phosphoethanolamine (TR-DHPE) (Molecular Probes, Eugene, OR) was first dissolved in chloroform and subsequently added to lipid spreading solutions at 0.5 mol%. All lipid solutions were stored at −20°C in glass vials. All water used was purified with a Milli-Q Ultrapure water purification system (Millipore, Bedford, MA).

### Surface activity and membrane insertion measurements

To evaluate the surface activity of the tau proteins, π measured by the adsorption of the proteins from a water subphase to an air/water interface was measured. The experiments were carried out at 25 °C using a MiniMicro Langmuir trough (KSV Instruments Ltd., Finland) with 45 mL of water as the subphase. A Wilhelmy plate sensor measured π of the air/subphase interface where π = γ_0_ − γ and γ_0_ is the surface tension of a clean air/water interface and γ is the surface tension of the air/water interface with adsorbed protein. The trough had a working surface area of 86.39 cm^2^. Before injecting protein into the subphase, barriers were partially closed to give a total surface area of 45 cm^2^. A gastight glass microsyringe (Hamilton, Reno, NV) was used to inject 1 mL of 45 μM tau to achieve a final tau concentration of 1 μM in the subphase.

To evaluate the interaction between tau and membranes, the insertion of tau into lipid monolayers at the air/water interface held at 25 mN/m was measured in a Langmuir trough as previously described^69^. All experiments were carried out on water subphase and 25 °C. All experiments were carried out at 25 mN/m and 1 μM tau concentration in the subphase. 1 mL of 45 μM tau in was then injected into the subphase underneath the monolayer and allowed to equilibrate with the lipid monolayer. Tau concentration in the subphase was 1 μM. Favorable tau−lipid interactions leading to the insertion of tau into the lipid monolayer caused an expansion of the monolayer surface area. The percent area expansion, or the change in the effective area per lipid molecule, is defined as % ΔA/A = 100(A − A_i_)/A_i_, where A is the surface area at time t and A_i_ is the initial surface area of the monolayer when prior to the injection of the protein at 25 mN/m. Videos of the protein-lipid film at the air/water interface was taken with a Nexus 5X model LG-H790 smartphone.

### Fluorescence Microscopy

To visualize lipid monolayer morphological change during tau insertion, the Langmuir trough was positioned on top of a motorized stage of an inverted fluorescence microscope (Olympus IX 71) with a 50x objective centered on a quartz window in the bottom of the trough. A 100 Watt mercury lamp was used for fluorescence excitation. Fluorescence images were collected by a QImaging camera (EXi Blue, QImaging Photometrics) and analyzed using the software QCapture Pro. TR-DHPE (0.5 mol%) was included in the spreading solution.

### GIXD measurements

GIXD measurements were collected at the BW1 beamline (HASYLAB, DESY, Hamburg, Germany) for tau insertion into DMPG monolayers. The synchrotron X-ray beam was monochromated to a wavelength (*λ*) of 1.30 Å by Bragg reflection from a Beryllium (200) monochromator crystal in Laue geometry. All experiments were carried out in an ultra-small volume Langmuir trough liquid diffractometer (20 mL subphase volume) at 23 °C and 1 μM tau concentration in water. The trough was temperature controlled and equipped with a Wilhelmy balance for surface pressure measurements and a motorized barrier.

The X-ray beam strikes the surface at an incident angle of 0.11°, which corresponds to a *Q*_z_ = 0.85*Q*_c_, where *Q*_c_ = 0.02176 Å^−1^ is the critical scattering vector for total external reflection. At this angle, the incident wave is totally reflected, while the refracted wave becomes evanescent with penetration depth of approximately 100 Å which maximizes surface sensitivity. The X-ray beam footprint on the liquid surface was approximately 2 × 50 mm^2^, bigger than the width of the ultra-small Langmuir trough used. This caused over illumination of the sample and small increases in the scattering background. The scattered intensity was measured by scanning over a range of horizontal scattering vectors, *Q*_xy_^54, 70^. Bragg peaks are intensity resolved in the *Q*_xy_-direction and integrated over channels along the z-direction (normal to the liquid subphase) in the 1-D position sensitive detector. The position of the maxima of the Bragg peaks, *Q*_xy_^max^, was used to calculate the repeat distances *d* = 2π/*Q*_xy_ of the 2-D lattice. The widths of the peaks, corrected for the instrument resolution, were used to determine the 2-D crystalline in-plane coherence length, *L*_xy_. The intensities resolved in *Q_z_* direction but integrated over the *Q_xy_* range of the Bragg peaks, so called Bragg rods, were also recorded and analyzed. The analysis of the Bragg rods provides the length (*L*_c_) of the coherently scattering moieties participating in the Bragg reflection. For lipid scattering, *L*_c_ is the length of the coherently scattering portions of the alkyl tails measured along their backbones. For protein β-sheet scattering, *L*_c_ is the length of the molecular moiety participating in coherent scattering.

## Data Availability

The data that support the findings are available from the corresponding author upon reasonable request.

## Acknowledgments

We gratefully acknowledge support from the National Science Foundation (Award Number 1150855), Alzheimer’s Association (NIRG-09-132478), UNM Research Allocation Committee, Oak Ridge Associated Universities Ralph E. Powe Junior Faculty Enhancement Award. E.M.J. acknowledges fellowship support from the NSF-IGERT program “Integrative Nanoscience and Microsystems” and the NCI Alliance for Nanotechnology in Cancer. C.M.V.Z. was supported by a postdoctoral fellowship from ASERT IRACDA K12 GM088021. This work benefited from the use the BW1 Beamline and Dr. Bernd Struth at the Deutsches Elektronen-Synchrotron in Hamburg, Germany. NSF provided support for JM to contribute to this project through the Independent Research and Development program. Any opinion, findings, and conclusions or recommendations expressed in this material are those of the author(s) and do not necessarily reflect the views of the National Science Foundation. E.M. and J.B. acknowledge support from the Max-Planck-Society (Max-Planck-Unit for Structural Molecular Biology at DESY, Hamburg) and the German Center for Neurodegenerative Diseases (DZNE, Bonn).

## Author Contributions

EYC, JM, and EM designed research; EMJ, JM, CVZ and JB performed research; EMJ and JM analyzed data; and EYC, JM, EMJ, and EM wrote the paper.

## Supplementary Information

**Figure S1:**
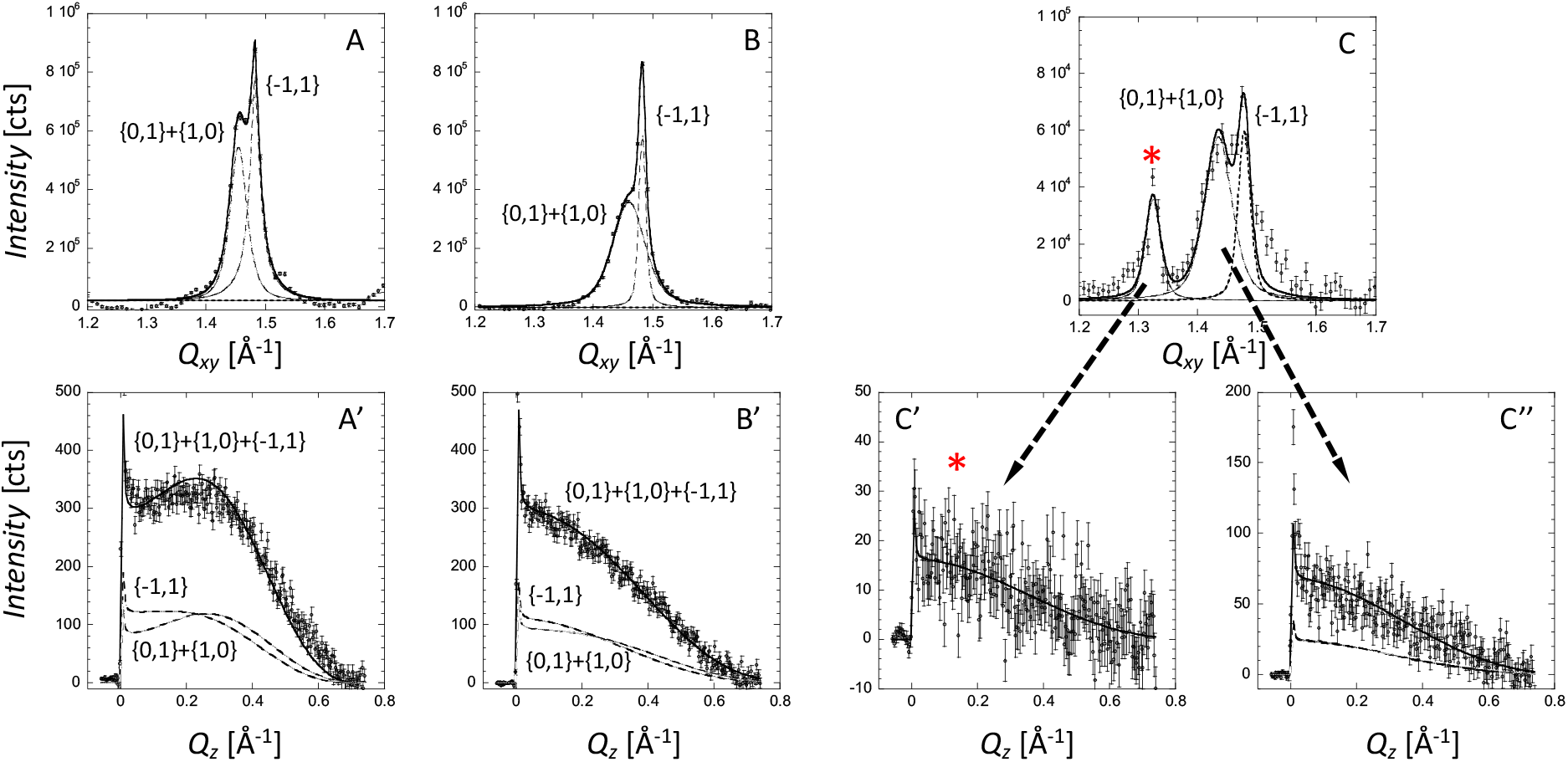
*Bragg peaks* (top row) and *rods* (bottom row) of a DMPG monolayer at the air/water interface at 25 mN/m and 25°C before (A, A’) and *t*_1_ = 2.5 hr (B, B’) and *t*_2_ = 12 hr (C, C’ and C”) after the injection of hTau40. *Bragg peaks* were obtained by integrating over the 0.05 Å^−1^ ≤ *Q_z_* ≤ 0.75 Å^−1^ region. The *Bragg peaks* (A, B, C) were fitted using the sum of Voigt profiles (solid line) and de-convoluted into separate peaks (dashed lines) corresponding to the two {1,0}+{0,1} and {1,−1} Bragg peaks. A’, B’, and C” show the sum of the two DMPG {1,0}+{0,1} and {1,−1} *Bragg rods* at times *t*_1_ and *t*_2_ after the injection of hTau40. The *Bragg rods* were obtained by integrating over the 1.35 Å^−1^ ≤ *Q*_xy_ ≤ 1.55 Å^−1^ region and fitted (solid line) by approximating the coherently scattering part of the alkyl tail by a cylinder of constant electron density. Each of the {0,1}, {1,0} and {−1,1} *Bragg rods* are shown as dashed lines in bottom row. The *Bragg peaks* and *rods* associated with the tau protein is indicated by red asterisks *. The protein peak *Bragg rod* (C’) were obtained by integrating over the 1.25 Å^−1^ ≤ *Q*_xy_ ≤ 1.35 Å^−1^ region.

**Figure S2:**
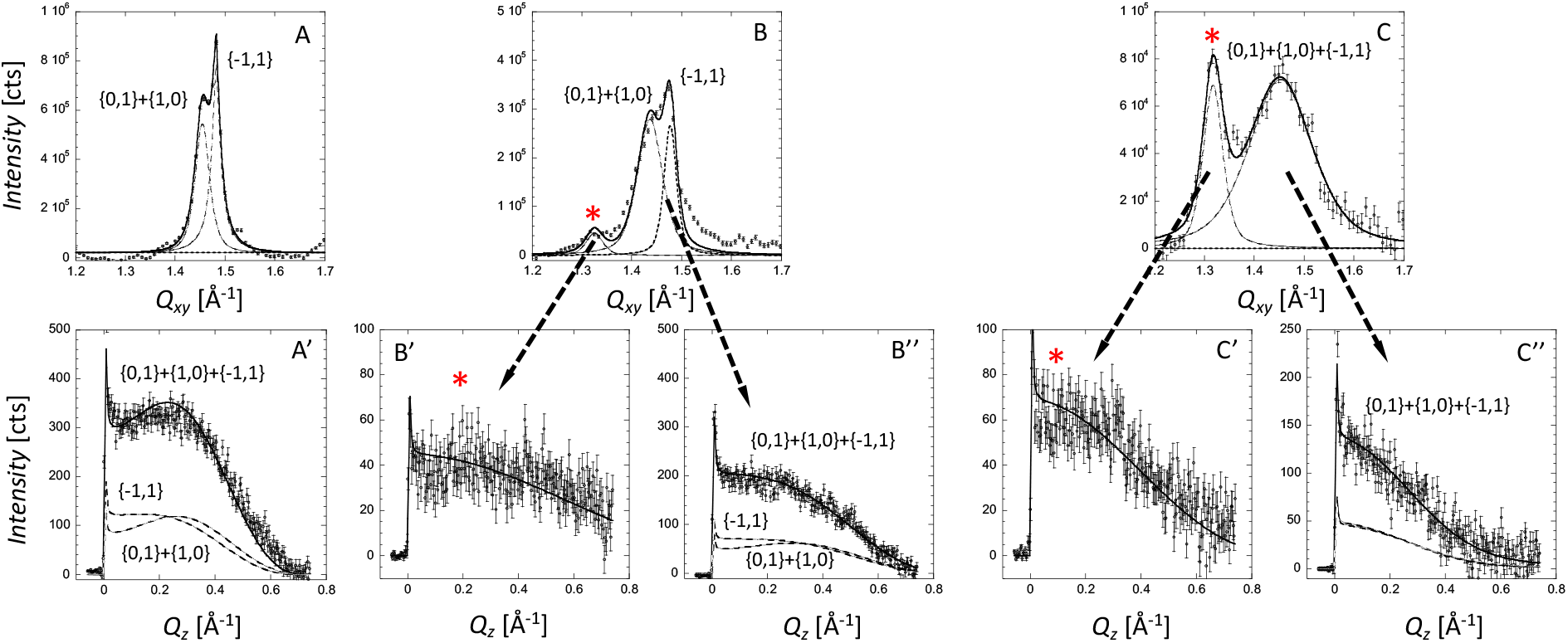
*Bragg peaks* (top row) and *rods* (bottom row) of a DMPG monolayer at the air/water interface at 25 mN/m and 25°C before (A, A’) and *t*_1_ = 2.5 hr (B, B’) and *t*_2_ = 12 hr (C, C’ and C”) after the injection of K18. *Bragg peaks* were obtained by integrating over the 0.05 Å^−1^ ≤ *Q_z_* ≤ 0.75 Å^−1^ region. The *Bragg peaks* were fitted using the sum of two Voigt profiles (solid line) and de-convoluted into separate peaks (dashed lines) corresponding to the two {1,0}+{0,1} and {1,−1} Bragg peaks. A’, B”, and C” show the sum of the two DMPG {1,0}+{0,1} and {1,−1} *Bragg rods t*_1_ and *t*_2_ after the injection of K18. The *Bragg rods* were obtained by integrating over the 1.35 Å^−1^ ≤ *Q*_xy_ ≤ 1.55 Å^−1^ region and fitted (solid line) by approximating the coherently scattering part of the alkyl tail by a cylinder of constant electron density. Each of the {0,1}, {1,0} and {−1,1} *Bragg rods* are shown as dashed lines in bottom row. The *Bragg peaks* and *rods* associated with the tau protein is indicated by *. The protein peak *Bragg rod* (B’ and C’) were obtained by integrating over the 1.25 Å^−1^ ≤ *Q*_xy_ ≤ 1.35 Å^−1^ region.

**Figure S3:**
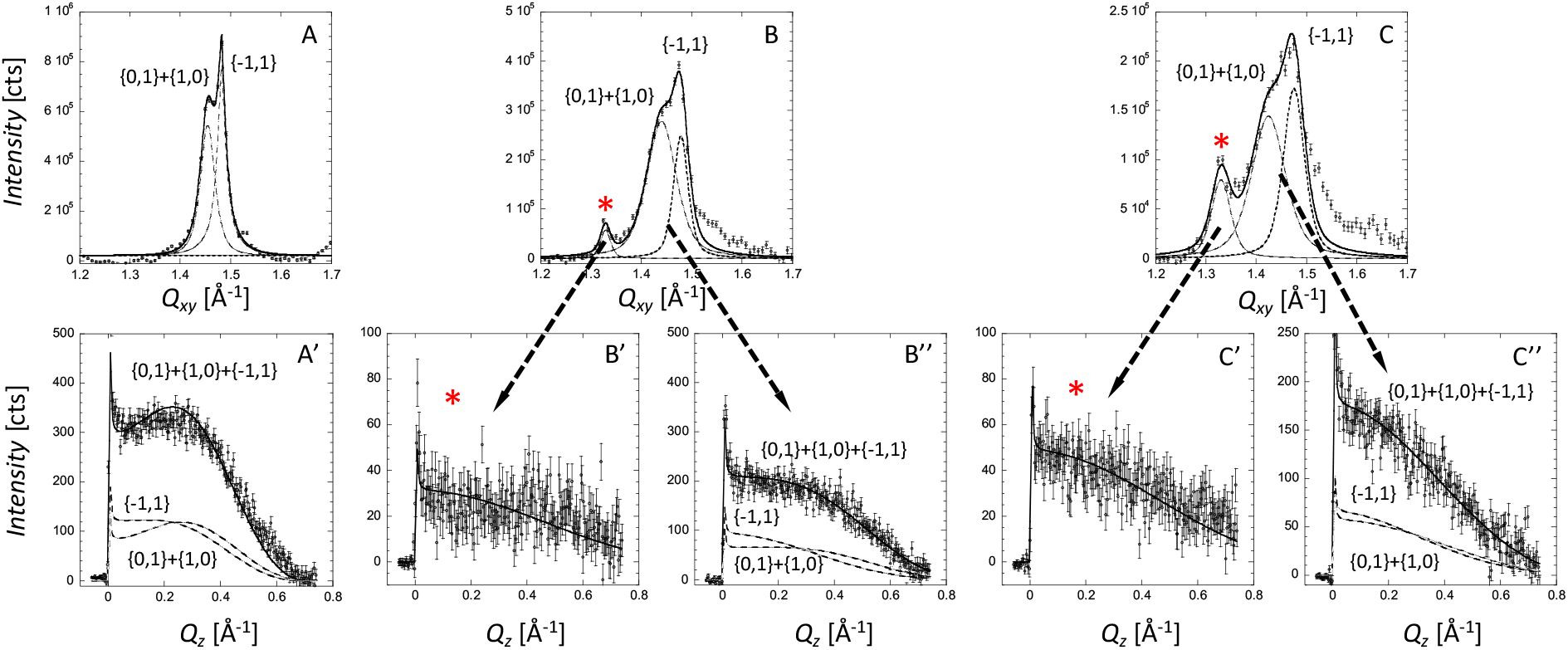
*Bragg peaks* (top row) and *rods* (bottom row) of a DMPG monolayer at the air/water interface at 25 mN/m and 25°C before (A, A’) and *t*_1_ = 2.5 hr (B, B’) and *t*_2_ = 12 hr (C, C’ and C”) after the injection of hTau40/3Epi. *Bragg peaks* were obtained by integrating over the 0.05 Å^−1^ ≤ *Q_z_* ≤ 0.75 Å^−1^ region. The *Bragg peaks* were fitted using the sum of two Voigt profiles (solid line) and de-convoluted into separate peaks (dashed lines) corresponding to the two {1,0}+{0,1} and {1,−1} Bragg peaks. A’, B”, and C” show the sum of the two DMPG {1,0}+{0,1} and {1,−1} *Bragg rods t*_1_ and *t*_2_ after the injection of hTau40/3Epi. The *Bragg rods* were obtained by integrating over the 1.35 Å^−1^ ≤ *Q*_xy_ ≤ 1.55 Å^−1^ region and fitted (solid line) by approximating the coherently scattering part of the alkyl tail by a cylinder of constant electron density. Each of the {0,1}, {1,0} and {−1,1} *Bragg rods* are shown as dashed lines in bottom row. The *Bragg peaks* and *rods* associated with the tau protein is indicated by red asterisks *. The protein peak *Bragg rod* (B’ and C’) were obtained by integrating over the 1.25 Å^−1^ ≤ *Q*_xy_ ≤ 1.35 Å^−1^ region.

**Table S1.** Structural parameters obtained from GIXD measurements of the full-length wildtype hTau40:DMPG film at the air/water interface at 25°C before (DMPG) and at two time points (*t*_1_ = 2.5 hrs and *t*_2_ = 12 hrs) after the injection of hTau40.

**Table S1A:** Structural parameters extracted from the DMPG diffraction peaks.

| Sample<br>(Surface Pressure) | Distorted hexagonal unit cell dimensions<br>$a, b, \gamma$<br>(Å, Å, degrees) | | | Area per lipid molecule<br>(Å<sup>2</sup>) | Integrated intensity<br>(%) | Coherence length<br>$L_c$<br>(Å) | Tilt angle<br>$t$<br>(°) | Tilt dir. from NN, non-symmetry<br>(°) | $\sigma$<br>(Å) |
| --- | --- | --- | --- | --- | --- | --- | --- | --- | --- |
| DMPG<br>(25 mN/m) | 4.93<br>$\pm 0.01$ | 4.93<br>$\pm 0.01$ | 118.8<br>$\pm 0.4$ | 42.6<br>$\pm 0.1$ | 46073<br>(100) | 14.0<br>$\pm 0.5$ | 18.1<br>$\pm 1.0$ | 0 | 0.5<br>$\pm 0.2$ |
| hTau40:DMPG<br>$t_1 = 2.5$ hrs<br>(34 mN/m) | 4.92<br>$\pm 0.01$ | 4.92<br>$\pm 0.01$ | 119.0<br>$\pm 0.4$ | 42.4<br>$\pm 0.2$ | 41697<br>(90) | 11.5<br>$\pm 0.5$ | 15.5<br>$\pm 1.0$ | 0 | 0.6<br>$\pm 0.2$ |
| hTau40:DMPG<br>$t_2 = 12$ hrs<br>(39 mN/m) | 4.96<br>$\pm 0.01$ | 4.96<br>$\pm 0.01$ | 118.0<br>$\pm 0.4$ | 43.4<br>$\pm 0.1$ | 6143<br>(13) | 11.7<br>$\pm 0.5$ | 16.7<br>$\pm 1.0$ | 0 | 0.6<br>$\pm 0.2$ |

Due to the constant area of the trough the protein insertion into the lipid monolayer following its injection caused the surface pressure of the protein/lipid film to increase during the experiment.
$L_c$ is the length of the coherently scattering part of the alkyl tail measured along its backbone.
Tilt angle is measured from the surface normal. The tilt angle is measured between the direction of nearest neighbor and the projection of the alkyl tail on the subphase surface. Nearest neighbor (NN) is along $a + b$ , where $a$ and $b$ are the 2D unit cell vectors. $\sigma$ is the Debye-Waller factor or root mean-square molecular displacement.

**Table S1B:** In-plane coherence length (*L*_*xy*_) values of the DMPG diffraction peaks.

| Sample<br>(Surface Pressure) | In-plane Bragg peaks coherence length<br>$L_{xy}$ (Å) $\pm 10$ Å | |
| --- | --- | --- |
| | $L_{10+01}$ | $L_{1-1}$ |
| DMPG | 170 | 310 |
| hTau40:DMPG $t_1 = 2.5$ hrs | 90 | 500 |
| hTau40:DMPG $t_2 = 12$ hrs | 100 | 240 |

$L_{xy}$ is the in-plane coherence length, an average length over which the scattering units are in registry.

**Table S1C:** Structural parameters extracted from the hTau40 protein peak at *Q*_*xy*_=1.34Å^−1^.

| Sample<br>(Surface Pressure) | $d$-spacing<br>(Å) | Integrated intensity<br>(%) | Coherence length<br>$L_c$ (Å) | Tilt angle<br>$t$ (°) | $\sigma$<br>(Å) | In-plane Bragg peaks coherence length<br>$L_{xy}$ (Å) |
| --- | --- | --- | --- | --- | --- | --- |
| hTau40<br>$t_1 = 2.5$ hrs | - | - | - | - | - | - |
| hTau40<br>$t_2 = 12$ hrs | $4.74 \pm 0.02$ | 1397<br>(100) | $7.2 \pm 1$ | 0 | $0.83 \pm 0.2$ | $200 \pm 10$ Å |

$L_c$ is the length of the coherently scattering part of the protein.
$L_{xy}$ is the in-plane coherence length, an average length over which the scattering units are in registry.
$\sigma$ is the Debye-Waller factor or root mean-square molecular displacement.

**Table S2.** Structural parameters obtained from GIXD measurements of truncated tau construct K18:DMPG film at the air/water interface at 25°C before (DMPG) and at two time points after (*t*_1_ = 2.5 hr and *t*_2_ ~12 hr) the injection of K18.

**Table S2A:** Structural parameters extracted from the DMPG diffraction peaks.

| Sample<br>(Surface Pressure) | Distorted hexagonal unit cell dimensions<br>$a, b, \gamma$<br>(Å, Å, degrees) | | | Area per lipid molecule<br>(Å<sup>2</sup>) | Integrated intensity (%) | Coherence length<br>$L_c$<br>(Å) | Tilt angle<br>$t$<br>(°) | Tilt dir. from NN, non-symmetry (°) | $\sigma$<br>(Å) |
| --- | --- | --- | --- | --- | --- | --- | --- | --- | --- |
| DMPG<br>(25 mN/m) | 4.93<br>$\pm 0.01$ | 4.93<br>$\pm 0.01$ | 118.8<br>$\pm 0.4$ | 42.6<br>$\pm 0.1$ | 46073<br>(100) | 14.0<br>$\pm 0.5$ | 18.1<br>$\pm 1.0$ | 0 | 0.5<br>$\pm 0.2$ |
| K18:DMPG<br>$t_1 = 2.5$ hrs<br>(45 mN/m) | 4.96<br>$\pm 0.01$ | 4.96<br>$\pm 0.01$ | 118.1<br>$\pm 0.4$ | 43.4<br>$\pm 0.2$ | 32586<br>(70) | 11.6<br>$\pm 0.5$ | 19.5<br>$\pm 1.0$ | 0 | 0.5<br>$\pm 0.2$ |
| K18:DMPG<br>$t_2 = 12$ hrs<br>(46 mN/m) | 5.00<br>$\pm 0.01$ | 5.00<br>$\pm 0.01$ | 120 | 43.2<br>$\pm 0.2$ | 14327<br>(30) | 10.5<br>$\pm 0.5$ | 13.6<br>$\pm 1.0$ | 0 | 0.6<br>$\pm 0.2$ |

Due to the constant area of the trough the protein insertion into the lipid monolayer following its injection caused the surface pressure of the protein/lipid film to increase during the experiment.
$L_c$ is the length of the coherently scattering part of the alkyl tail measured along its backbone.
Tilt angle is measured from the surface normal. The tilt angle is measured between the direction of nearest neighbor and the projection of the alkyl tail on the subphase surface. Nearest neighbor (NN) is along $a + b$ , where $a$ and $b$ are the 2D unit cell vectors. $\sigma$ is the Debye-Waller factor or root mean-square molecular displacement.

**Table S2B:** In-plane coherence length (*L_xy_*) values of the DMPG diffraction peaks.

| Sample<br>(Surface Pressure) | In-plane Bragg peaks coherence length<br>$L_{xy}$ (Å) $\pm 10$ Å | |
| --- | --- | --- |
| | $L_{10+01}$ | $L_{1-1}$ |
| DMPG | 170 | 310 |
| K18:DMPG, $t_1 = 2.5$ hrs | 100 | 200 |
| K18:DMPG, $t_2 = 12$ hrs | 40 | 40 |

$L_{xy}$ is the in-plane coherence length, an average length over which the scattering units are in registry.

**Table S2C:** Structural parameters extracted from the K18 protein peak at *Q_xy_*=1.34Å^−1^.

| Sample<br>(Surface Pressure) | $d$-spacing<br>(Å) | Integrated intensity (%) | Coherence length<br>$L_c$ (Å) | Tilt angle<br>$t$ (°) | $\sigma$<br>(Å) | In-plane Bragg peaks coherence length<br>$L_{xy}$ (Å) |
| --- | --- | --- | --- | --- | --- | --- |
| K18<br>$t_1 = 2.5$ hrs | $4.74 \pm 0.03$ | 2415<br>(56) | 4.5* | 0 | $0.6 \pm 0.2$ | $150 \pm 20$ Å |
| K18<br>$t_2 = 12$ hrs | $4.77 \pm 0.02$ | 4272<br>(100) | $6.5 \pm 1$ | 0 | $0.7 \pm 0.2$ | $120 \pm 10$ Å |

$L_c$ is the length of the coherently scattering part of the protein.
$L_{xy}$ is the in-plane coherence length, an average length over which the scattering units are in registry.
$\sigma$ is the Debye-Waller factor or root mean-square molecular displacement.
\* Due to weak scattering and high background, the extraction of the *Bragg rod* and resulting structural data was with a higher degree of uncertainty.

**Table S3.** Structural parameters obtained from GIXD measurements of the hyperp-hosphorylation mimic mutant hTau40/3Epi:DMPG film at the air/water interface at 25°C before (DMPG) and at two time points after (*t*_1_ = 2.5 hr and *t*_2_ ~12 (8.5?) hr) the injection of hTau40/3EPi.

**Table S3A:** Structural parameters extracted from the DMPG diffraction peaks.

| Sample<br>(Surface Pressure) | Distorted hexagonal unit cell dimensions<br>$a, b, \gamma$<br>(Å, Å, degrees) | | | Area per lipid molecule<br>(Å<sup>2</sup>) | Integrated intensity (%) | Coherence length<br>$L_c$<br>(Å) | Tilt angle<br>$t$<br>(°) | Tilt dir. from NN, non-symmetry (°) | $\sigma$<br>(Å) |
| --- | --- | --- | --- | --- | --- | --- | --- | --- | --- |
| DMPG<br>(25 mN/m) | 4.93<br>$\pm 0.01$ | 4.93<br>$\pm 0.01$ | 118.8<br>$\pm 0.4$ | 42.6<br>$\pm 0.1$ | 46073<br>(100) | 14.0<br>$\pm 0.5$ | 18.1<br>$\pm 1.0$ | 0 | 0.5<br>$\pm 0.2$ |
| hTau40/3Epi:<br>DMPG<br>$t_1 = 2.5$ hrs<br>(47 mN/m) | 4.95<br>$\pm 0.01$ | 4.95<br>$\pm 0.01$ | 118.2<br>$\pm 0.4$ | 43.2<br>$\pm 0.1$ | 37070<br>(80) | 10.9<br>$\pm 0.5$ | 18.0<br>$\pm 1.0$ | 0 | 0.6<br>$\pm 0.1$ |
| hTau40/3Epi:<br>DMPG<br>$t_2 = 12$ hrs<br>(44 mN/m) | 4.98<br>$\pm 0.01$ | 4.98<br>$\pm 0.01$ | 117.6<br>$\pm 0.4$ | 43.9<br>$\pm 0.1$ | 26549<br>(58) | 10.9<br>$\pm 0.5$ | 19.2<br>$\pm 1.0$ | 0 | 0.6<br>$\pm 0.2$ |

Due to the constant area of the trough the protein insertion into the lipid monolayer following its injection caused the surface pressure of the protein/lipid film to increase during the experiment.
$L_c$ is the length of the coherently scattering part of the alkyl tail measured along its backbone.
Tilt angle is measured from the surface normal. The tilt angle is measured between the direction of nearest neighbor and the projection of the alkyl tail on the subphase surface. Nearest neighbor (NN) is along $a + b$ , where $a$ and $b$ are the 2D unit cell vectors. $\sigma$ is the Debye-Waller factor or root mean-square molecular displacement.

**Table S3B:**
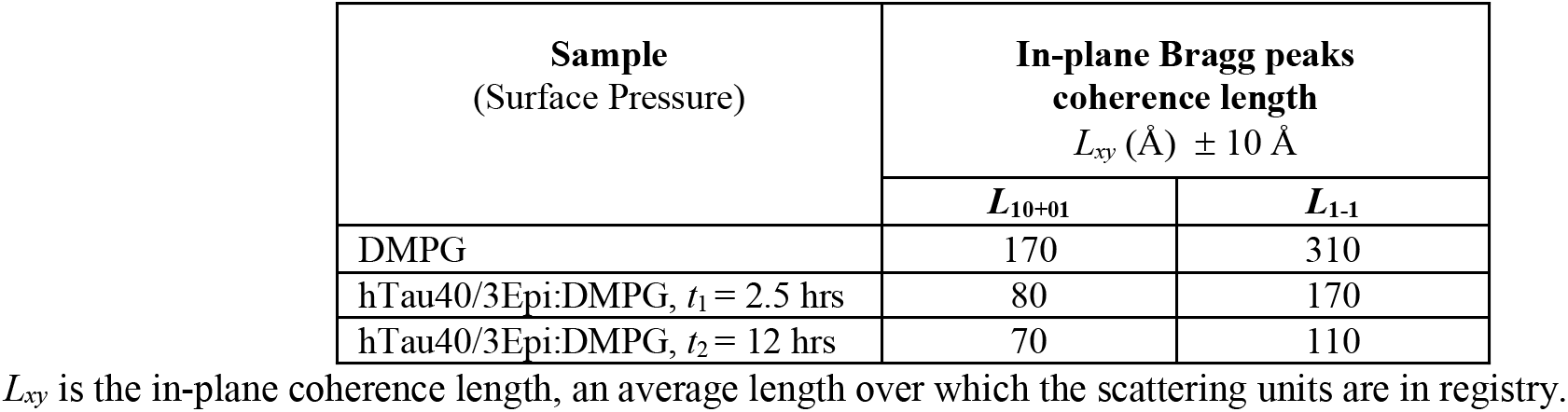
In-plane coherence length (*L_xy_*) values of the DMPG diffraction peaks.

**Table S3C:** Structural parameters extracted from the hTau40/3Epi protein peak at *Q_xy_*=1.34Å^−1^.

| Sample<br>(Surface Pressure) | $d$-spacing<br>(Å) | Integrated intensity (%) | Coherence length<br>$L_c$ (Å) | Tilt angle<br>$t$ (°) | $\sigma$<br>(Å) | In-plane Bragg peaks coherence length<br>$L_{xy}$ (Å) |
| --- | --- | --- | --- | --- | --- | --- |
| hTau40/3Epi<br>$t_1 = 2.5$ hrs | 4.72<br>$\pm 0.02$ | 1789<br>(37) | 5.5<br>$\pm 1$ | 0 | 0.63<br>$\pm 0.2$ | 110 |
| hTau40/3Epi<br>$t_2 = 12$ hrs | 4.72<br>$\pm 0.02$ | 4795<br>(100) | 5.5<br>$\pm 1$ | 0 | 0.67<br>$\pm 0.2$ | 130 |

$L_c$ is the length of the coherently scattering part of the protein.
$L_{xy}$ is the in-plane coherence length, an average length over which the scattering units are in registry.
$\sigma$ is the Debye-Waller factor or root mean-square molecular displacement.

**Figure S4:**
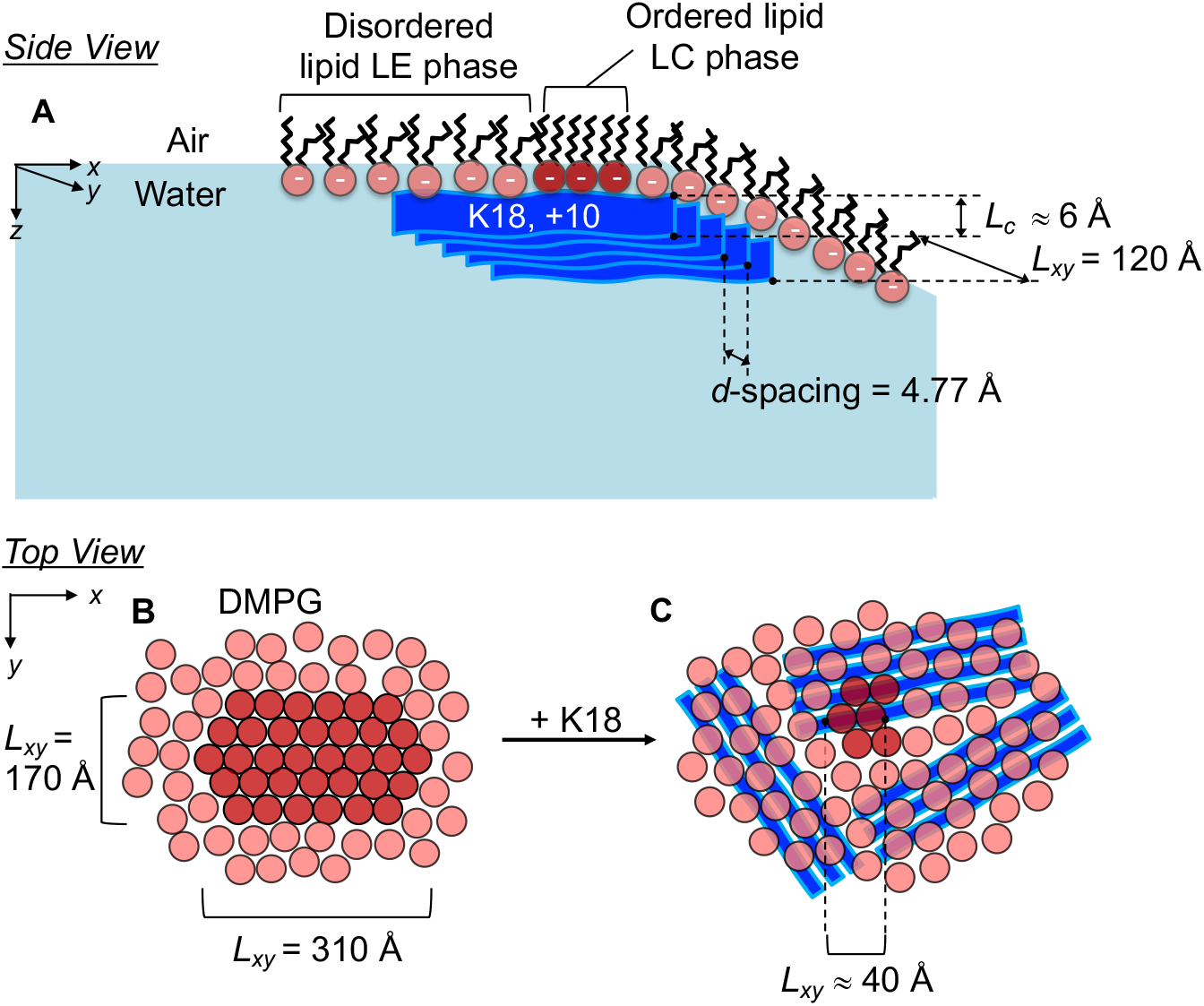
Schematics of β-sheet enriched K18 assemblies at the anionic DMPG monolayer surface. **A**: side view (parallel or in-plane to the lipid membrane) shows an ordered K18 assembly associated with the anionic DMPG monolayer at the air/water interface. *d*-spacing and coherence lengths (*L_xy_*) shown are for the K18 oligomer. *L_c_* is the out-of-plane length of the coherently scattering moiety of the K18 assembly. **B**: top views (perpendicular or out-of-plane to the lipid membrane) of the membrane before (**B**) and after (**C**) K18 association. The *L_xy_* values in these schematics are those of the ordered lipid domains. After 12 hrs the average size of the ordered DMPG domains deceased from about 300 Å to 40 Å (Table S3B). The orientation of the K18 β-sheet arrangements vis-à-vis the orientation of the DMPG lattice cannot be determined from the experimental data.

**Figure S5:**
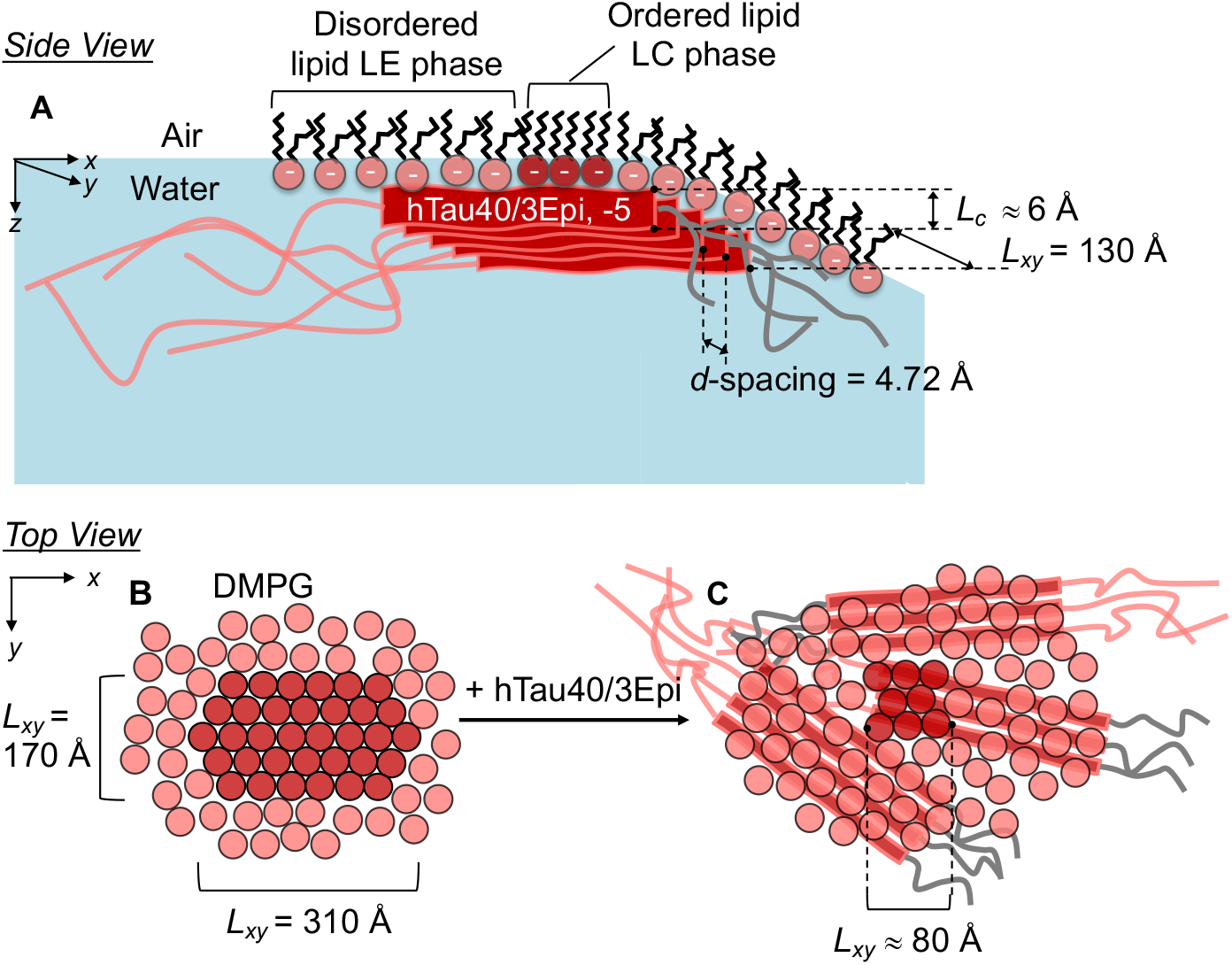
Schematics of β-sheet enriched hTau40/3Epi assemblies at the anionic DMPG monolayer surface. **A**: side view (parallel or in-plane to the lipid membrane) shows an ordered hTau40/3Epi assembly associated with the anionic DMPG monolayer at the air/water interface. *d*-spacing and coherence lengths (*L_xy_*) shown are for the hTau40/3Epi oligomer. *L_c_* is the out-of-plane length of the coherently scattering moiety of the hTau40/3Epi assembly. B. top views (perpendicular or out-of-plane to the lipid membrane) of the membrane before (left) and after (right) hTau40/3Epi association. The *L_xy_* values in these schematics are those of the ordered lipid domains. After 12 hrs the average size of the ordered DMPG domains deceased from about 300 Å to 80 Å (Table S2B). The orientation of the hTau40/3Epi β-sheet arrangements vis-à-vis the orientation of the DMPG lattice cannot be determined from the experimental data.

---

**Video S1**: 1 μM hTau40 incubated for approximately 12 hours with a DMPG monolayers held at 25 mN/m on water subphase at 25°C. In the video, the platinum Wilhelmy plate is seen moving around and in and out of the water subphase containing the lipid-tau protein film at the air/water interface. Note that the tau-DMPG film appears viscous and can be dragged up with the Wilhelmy plate as it is pulled up from the film at the air/water interface. Video can be viewed at: https://www.dropbox.com/s/diviwz54vxofz62/Video%201%20gelation_on_surface_hTau40%20and%20DMPG%20overnight.mp4?dl=0

**Video S2**: 0.25 μM amyloid-β peptide (Aβ40) incubated for approximately 12 hrs with a DMPG monolayers held at 25 mN/m on water subphase at 25°C. In the video, the platinum Wilhelmy plate is seen moving around and in and out of the water subphase containing the lipid-Aβ protein film at the air/water interface. Notice that the lipid-Aβ film breaks away easily from the Wilhelmy plate as it is pulled up from the film at the air/water interface. Video can be viewed at: https://www.dropbox.com/s/4oduaqmnzbezcwg/Video%203%20no_gelation_Amyloid%20beta%20and%20DMPG%20overnight.mp4?dl=0

